# A novel mitogenome for *Euterpe precatoria* (Arecaceae)

**DOI:** 10.64898/2026.09.28.754896

**Authors:** Lucile Soler, Francine Alves Noguiera de Almeida, Alexandre Gibau de Lima, Mai-Britt Mosbech, Christian Tellgren-Roth, Christine D. Bacon

## Abstract

Mitochondrial genomes of angiosperms are unusual due to their large size, low gene content, and high structural variation. In the palm family (Arecaceae), mitochondrial architecture is highly dynamic, characterized by expansive sizes, fragmentation, and extensive recombination activity. This biological complexity presents a significant bioinformatic challenge, especially since only three complete palm mitochondrial genomes are publicly available to date. Here, we present the complete mitochondrial genome sequence for the Amazonian palm *Euterpe precatoria*, assembled de novo using PacBio HiFi long-read sequencing data. The assembly required extensive manual curation and graph tracing to resolve a complex recombination topology. The final resolved sequence spans 1,268,308 base pairs and is supported by a mean read coverage depth estimated at 746X. Annotation revealed a high degree of structural complexity, including 177 total gene features corresponding to 76 unique genes. This gene set includes 41 unique protein-coding genes (56 features), 32 unique tRNA genes (118 features), and three unique rRNA genes, reflecting the characteristic gene duplication and fragmentation documented in Arecaceae. The complete mitochondrial genome sequence is publicly available and provides a high-quality resource for understanding the mechanisms of plant mitochondrial genome expansion and structural evolution.

## BACKGROUND AND SUMMARY

Mitochondria are fundamental to plant development, fitness, and reproduction. Their contribution to energy production, metabolism, and cell homeostasis relies on the performance of their own genome, which makes them semi-autonomous^1^. Mitochondrial genome evolution in flowering plants is unusual in comparison to other lineages in the tree of life, due to their large size, exceptionally low mutation rate, and high levels of structural variation^2^.

The palm family (Arecaceae) is a prime example of complex mitochondrial evolution^3-5^. Prior to this study, only three complete mitochondrial genomes from the Arecaceae were publicly available: *Cocos nucifera* L. (Cocoseae; Arecoideae)^3^, *Phoenix dactylifera* L. (Phoeniceae; Coryphoideae)^4^, *and Syagrus coronata* (Mart.) Becc. (Cocoseae; Arecoideae)^5^. These mitogenomes are notable for their large size (>640k bps), structural complexity, and high levels of recombination. Genomic rearrangements and gene transfer have been identified from both the chloroplast and nuclear genomes to palm mitochondrial genomes^3,5^. This complexity underscores the need for additional sampling to better understand mitochondrial genome evolution within the family.

Here, we develop a complete mitochondrial genome for *Euterpe precatoria* Mart. (Euterpeae; Arecoideae). Not only is it a widely distributed species across Central and South America, but it is also the most abundant species in Amazonia, with over five billion individuals found across Brazil, Colombia, Ecuador, French Guiana, Guiana, Peru, Suriname, and Venezuela^6^. This critical role in forest structure also means that *E. precatoria* plays a disproportionate role in the global carbon cycle, assimilating and storing more carbon than other species in the region^7^. The species is also of high social, cultural, and economic importance. In local Amazonian communities, different parts of the palm are used for handicrafts, construction material, and medicinal uses^8^, and the fruit is an important source of subsistence and income^9^. The global market value of the fruit of *E. precatoria* and its congener *E. oleracea* is forecasted to grow to 3.6 billion USD by 2030^10^. The species, therefore, urgently requires genomic resources^11^, for the understanding of its evolution, for crop development, and for conservation.

The mitochondrial genome sequence of *Euterpe precatoria* was assembled *de novo* using PacBio high-fidelity (HiFi) long-read sequencing data. The biological complexity of palm mitochondria presented significant bioinformatic challenges, as these genomes defy automated assembly into a single, simple circular sequence. Because of the structural complexity and the presence of multiple recombination pathways, the assembly required extensive manual curation and validation. This was achieved by carefully examining and resolving the assembly graph using Bandage, a tool for visualizing assembly graphs, to ensure the full mitochondrial gene content and connectivity were accurately captured. The final assembly is strongly supported by high-coverage data, with a mean read coverage depth of 746X across the genome. The dataset comprises a complete, single sequence representing the mitochondrial master content spanning 1,268,308 base pairs (bp), with a mean GC content of 42.9%. This high-quality dataset provides a valuable resource for comparative genomics within the Arecaceae and will facilitate future studies on the mechanisms driving mitochondrial genome expansion and structural evolution in this diverse plant family.

## DATA COLLECTION AND SEQUENCING

### Sample Collection

Leaf samples of *Euterpe precatoria* were collected into silica gel to preserve material for DNA extraction. We sampled an individual grown in the live collection (#443) from the Botanical Garden of Rio de Janeiro, Brazil. We made an herbarium specimen of the individual (#788749), which is deposited in the RB herbarium at the garden.

### DNA Extraction and Quality Control

DNA was extracted from a total of 1.6 grams cryo-ground leaves using CTAB lysis (2% CTAB, 100 mM Tris-HCl ph8, 20 mM EDTA, 1.4 M NaCl, 1% PEG 8000) followed by chloroform extraction and precipitation using 0.7x volume isopropanol. The pellet was resuspended in G2 buffer (Qiagen Cat #1014636) and purified according to the Genomic tip 100G (Qiagen Cat #10223) protocol with elution in TE-buffer (10 mM Tris, 0.1 mM EDTA). Concentration was measured using the Qubit Broad Range kit (Thermo Fisher Scientific # Q32850), purity was checked by UV-absorbance, and fragment lengths determined using the Genomic DNA 165 kb Kit (Agilent # FP-1002-0275) on the Femto Pulse System.

### Library Preparation and Sequencing Platform

For the preparation of the sequencing Library, 26 µg of DNA were sheared to a size of 10500 bp on a Megaruptor 3 instrument (Diagenode, Seraing, Belgium). The SMRTbell library was prepared according to PacBio protocol of preparing multiplexed amplicon libraries using SMRTbell® prep kit 3.0 (Pacific Biosciences, Menlo Park, CA, USA). SMRTbells were sequenced on a Revio system according to the manufacturer’s instructions (Pacific Biosciences, Menlo Park, CA, USA).

## MITOCHONDRIAL GENOME ASSEMBLY AND CURATION

### Sequencing Data and Reference Mapping

The complete mitochondrial genome sequence for *Euterpe precatoria* was derived from PacBio HiFi long-read sequencing data (ENA Project Accession: PRJEB105104). To isolate mitochondrial-derived sequences and guide the subsequent *de novo* assembly, raw reads were initially mapped against two reference mitochondrial genomes: *Cocos nucifera* (NC_031696), *Phoenix dactylifera* (NC_016740). Read mapping was performed using minimap2 version 2.26^12^ with the PacBio HiFi reads, using the command: minimap2 -ax map-hifi [reference].fasta all_reads_palm.fastq.gz > [reference]_all_reads.sam.

### Read Preparation and De-duplication

Reads mapped to each reference were subsequently isolated and processed using SAMtools version 1.16.1^13^ to generate a high-quality, de-duplicated input for *de novo* assembly. The processing pipeline for each read set included: (1) converting SAM files to BAM format and retaining only mapped reads (SAMtools view -b -F 4); (2) sorting reads by name and correcting alignment tags (SAMtools collate and samtools fixmate -m); (3) sorting by position and marking/removing duplicates (SAMtools sort and samtools markdup -r); and (4) converting the cleaned, de-duplicated BAM files to FASTA format (SAMtools fasta) for input into the assembler. In total, 342,342 reads were used in the mitochondrial final assembly.

### De Novo Assembly and Curation

The two prepared read sets were independently assembled *de novo* using the Flye assembler version 2.9.1^14^, explicitly configured for PacBio HiFi data: flye --pacbio-hifi [reads].fa --o flye_[reference]_assembly. The assembly generated from the *C. nucifera*-mapped reads was selected for final resolution because it produced a simpler assembly graph. The resulting assembly graph was visualized using Bandage version 0.8.1^15^ (Fig. 1).

**Figure 1.**
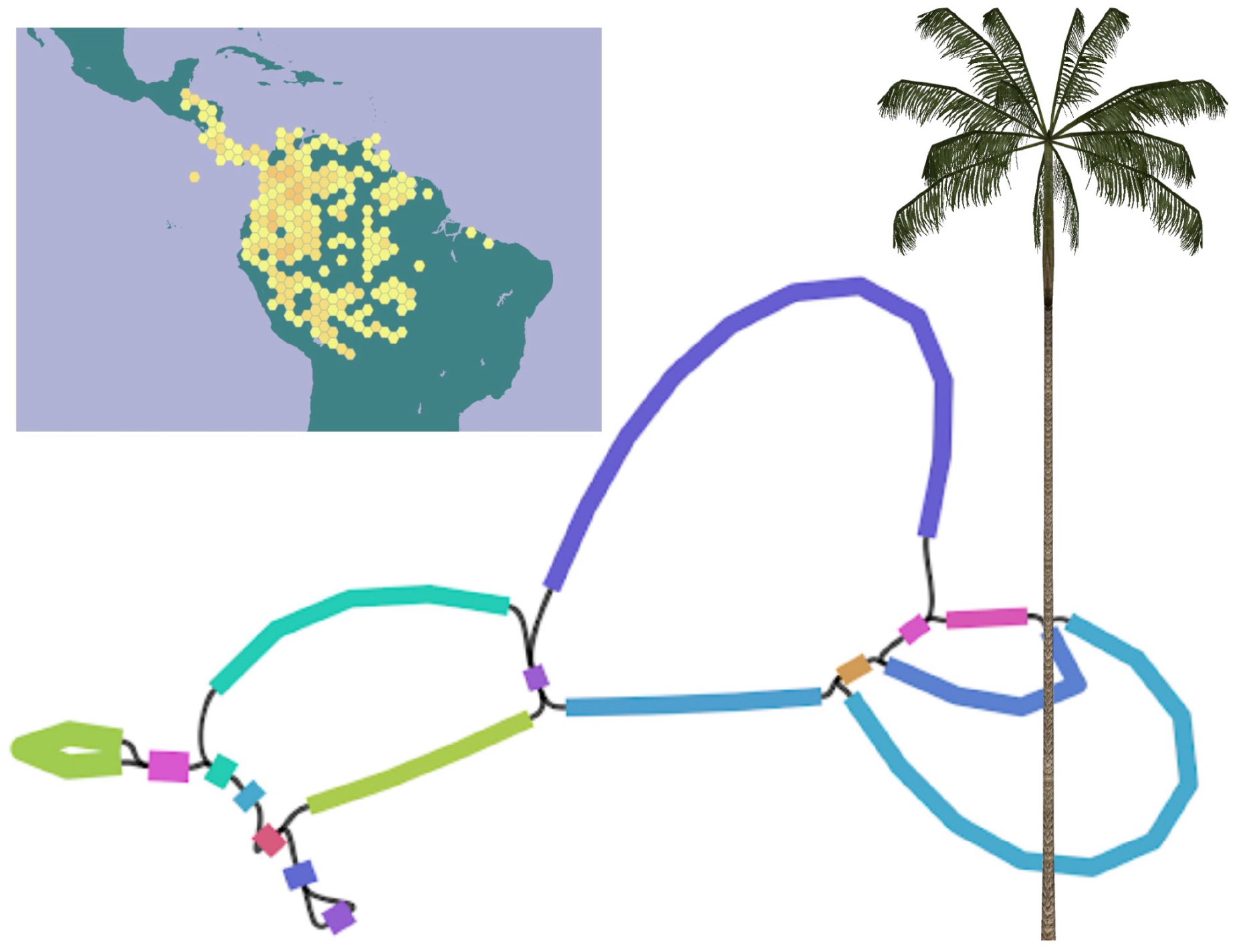
*Euterpe precatoria*, shown on the right, is a slender-stemmed, pinnate-leaved palm. It is a hyperdominant species in Amazonia with a wide distribution, as shown in the distribution map from the Global Biodiversity Information Facility (GBIF). Here, we developed genomic resources for the mitochondrial genome, here visualized in the assembly graph.

Initial automated attempts to achieve a simple, single-contig circularization failed, confirming that the assembly represents a complex, multi-path recombination structure. To obtain the final single, non-redundant sequence, the graph was resolved through manual tracing in Bandage. A persistent contaminating sequence, identified as a Nuclear Mitochondrial DNA segment (NUMT^16^) and corresponding to edge_4, was confirmed and intentionally excluded. The final master content sequence was constructed by concatenating the edges and orientations detailed in Supplementary Table S1. The final assembled mitochondrial genome sequence is 1,268,308 base pairs (bp) in length, with a mean GC content of 42.9%. Quality was validated by re-mapping all filtered reads against the final linear contig, confirming a uniform coverage of approximately 746 X and a mapping rate of 1.20%.

## MITOCHONDRIAL GENOME ANNOTATION AND RESULTS

### Gene and Feature Annotation Methods

The assembled sequence was annotated to identify functional elements. Annotation was performed using the integrated pipeline PMGA version 4^17^. PMGA employs a homology-based approach for identifying functional elements by comparing the assembled sequence against a comprehensive database of 319 known plant mitogenomes.

Transfer RNA (tRNA) genes were predicted using the tools integrated within PMGA: tRNAscan-SE 2.0^18^ and ARAGORN^19^, utilizing the plant mitochondrial genetic code. PMGA further incorporated CPGAVAS2^20^ for identifying potential plastid-derived genes, and tools like MISA^22^, Tandem Repeat Finder^22^, and Vmatch (REPuter)^23^ for identifying repeat sequences. Additionally, Deepred-MT^24^ was utilized to identify RNA editing sites in the coding sequences.

### Mitochondrial Gene Content

The complete mitochondrial genome annotation contained a total of 177 gene features, corresponding to 76 unique genes. This included 56 protein-coding gene features (41 unique genes) and the non-coding RNA components, reflecting a complex gene duplication profile common in Arecaceae^25^ (Table 1). Additionally, the complete annotation identified 118 tRNA genes (32 unique tRNAs) and 3 rRNA genes (3 unique rRNAs), consistent with a fully functional plant mitochondrial genome.

**Table 1.**
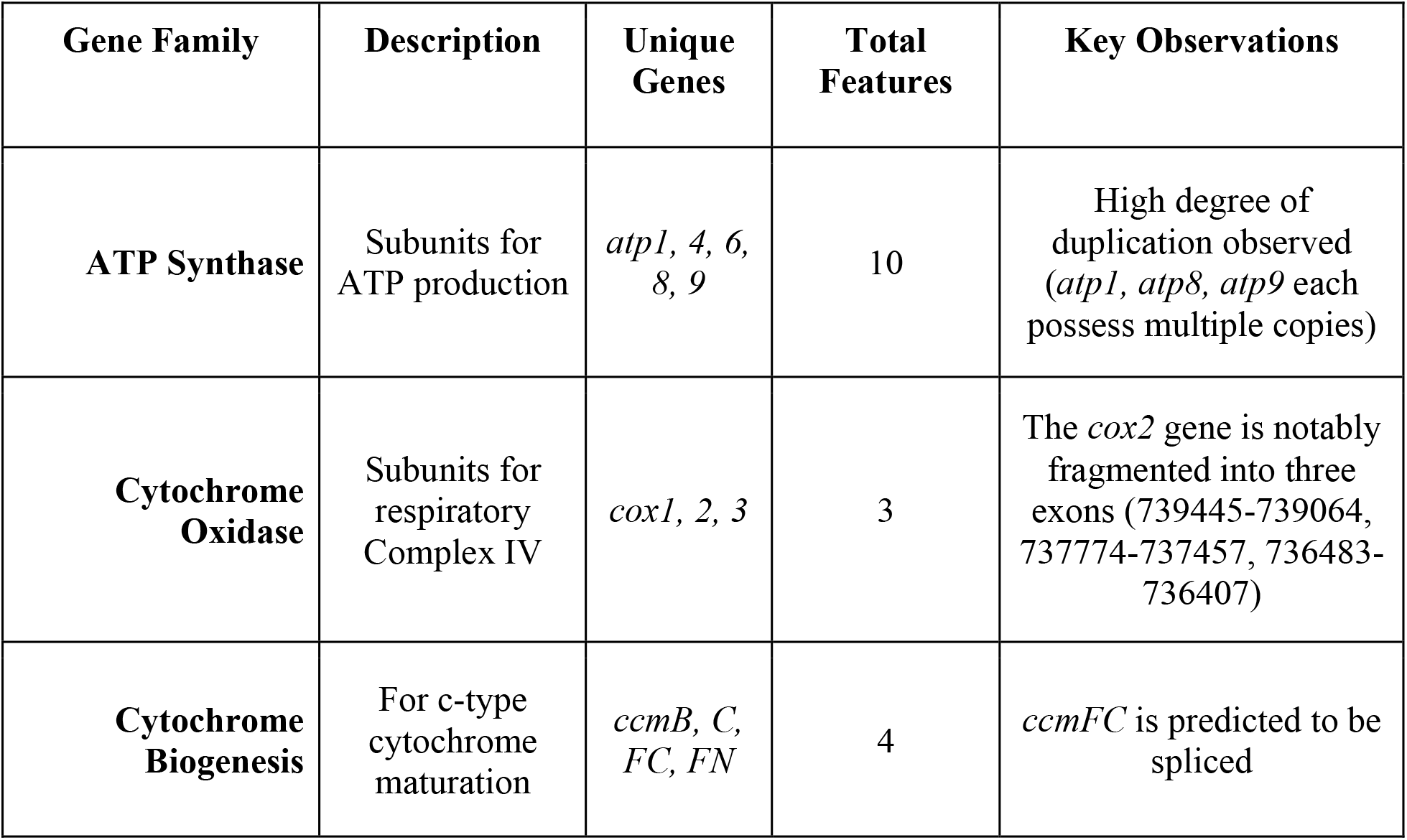
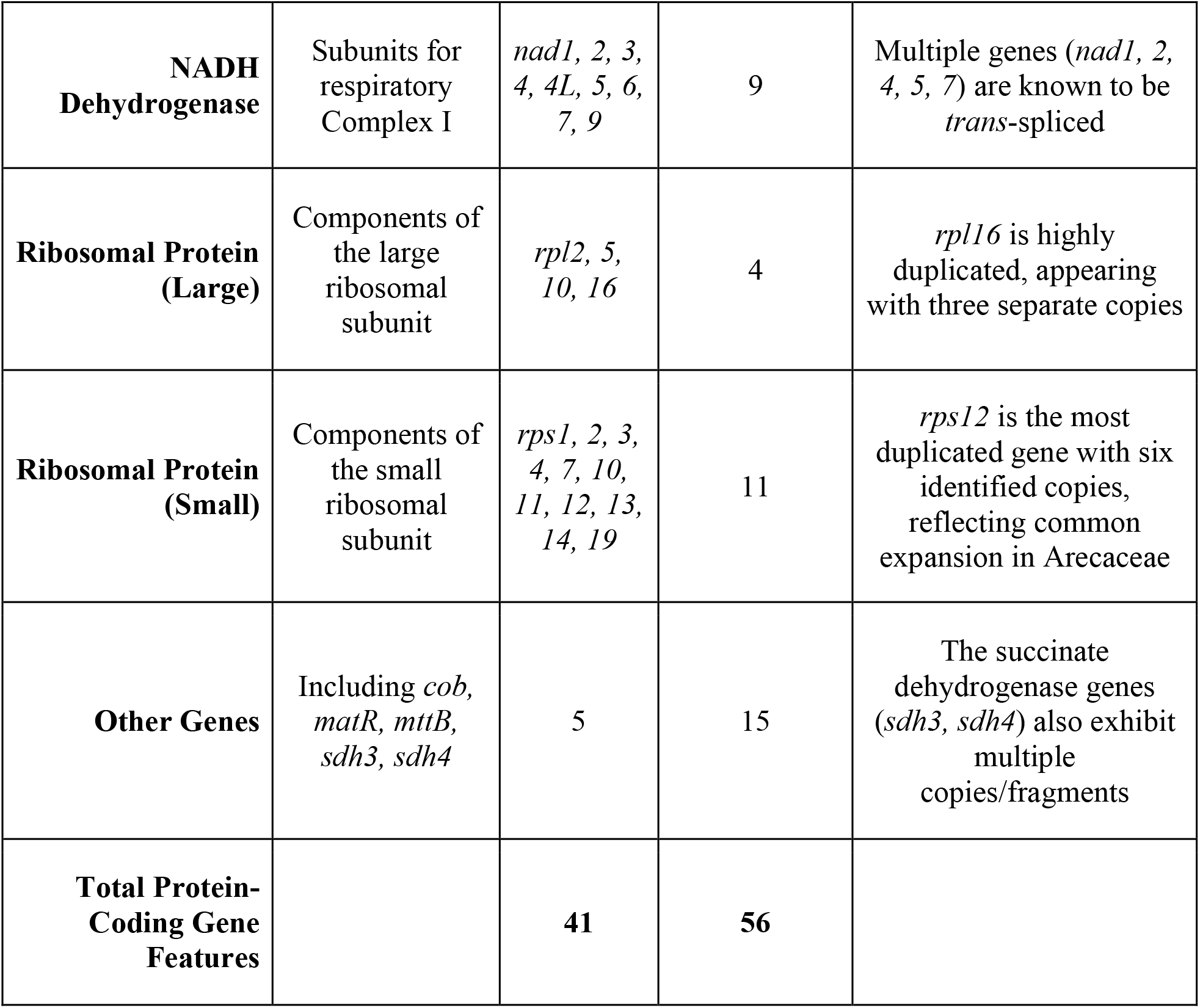
Results of the automated mitochondrial gene annotation using PMGA (Plant Mitochondrial Genome Annotator v4). The total features include copies.

| Gene Family | Description | Unique Genes | Total Features | Key Observations |
| --- | --- | --- | --- | --- |
| ATP Synthase | Subunits for ATP production | *atp1*, 4, 6, 8, 9 | 10 | High degree of duplication observed (*atp1*, *atp8*, *atp9* each possess multiple copies) |
| Cytochrome Oxidase | Subunits for respiratory Complex IV | *cox1*, 2, 3 | 3 | The *cox2* gene is notably fragmented into three exons (739445-739064, 737774-737457, 736483-736407) |
| Cytochrome Biogenesis | For c-type cytochrome maturation | *ccmB*, *C*, *FC*, *FN* | 4 | *ccmFC* is predicted to be spliced |
| **NADH Dehydrogenase** | Subunits for respiratory Complex I | *nad1*, 2, 3, 4, 4*L*, 5, 6, 7, 9 | 9 | Multiple genes (*nad1*, 2, 4, 5, 7) are known to be *trans*-spliced |
| **Ribosomal Protein (Large)** | Components of the large ribosomal subunit | *rpl2*, 5, 10, 16 | 4 | *rpl16* is highly duplicated, appearing with three separate copies |
| **Ribosomal Protein (Small)** | Components of the small ribosomal subunit | *rps1*, 2, 3, 4, 7, 10, 11, 12, 13, 14, 19 | 11 | *rps12* is the most duplicated gene with six identified copies, reflecting common expansion in Arecaceae |
| **Other Genes** | Including *cob*, *matR*, *mttB*, *sdh3*, *sdh4* | 5 | 15 | The succinate dehydrogenase genes (*sdh3*, *sdh4*) also exhibit multiple copies/fragments |
| **Total Protein-Coding Gene Features** |  | **41** | **56** |  |

## DATA RECORDS

The mitochondrial genome assembly and annotation are available as an SRA/ENA project at the accession PRJEB105104.

## TECHNICAL VALIDATION

To validate the 1,268,308 bp *Euterpe precatoria* mitochondrial assembly, we mapped the original PacBio HiFi reads to the final curated sequence using minimap2. The resulting alignment was processed using SAMtools to generate sorted and indexed BAM files. We calculated the mean coverage depth across the genome by applying an awk script to the output of samtools depth (awk ‘{sum+=$3; count++} END {print sum/count}’ mt_depth.txt), which resulted in a mean of 746X. Alignment statistics generated via samtools flagstat indicated that 1.20% of the total reads mapped to the mitochondrial sequence. Structural validation was performed by manually tracing the assembly graph in Bandage to resolve recombination pathways and exclude a specific nuclear mitochondrial DNA (NUMT) segment, identified as edge_4. The biological consistency of the assembly was assessed through homology-based annotation using the PMGA pipeline, which identified 76 unique genes. This gene content profile is consistent with previously documented mitochondrial genomes in the Arecaceae family.

## CODE AVAILABILITY

All software and pipelines are publicly accessible at https://github.com/NBISweden/C_BACON_Euterpe-precatoria_Mitochondria/tree/main.

## ACKNOWLEDGEMENTS

We acknowledge the Brazilian Ministry of the Environment for approval of the SISGEN permit #R82D7F5 for access to the genetic resources and traditional knowledge of the specimen (#788749) deposited at the RB herbarium at the Botanical Garden of Rio de Janeiro, Brazil, as well as the garden’s permission to collect from the live collection (#443). We acknowledge Yvonne Kallberg of NBIS for support with the ENA project organization and submission.

## AUTHOR CONTRIBUTIONS

L.S. and C.D.B. conceived the study. F.A. and A.G.L. collected and documented the plant material, M.-B.M. and C.T.-R. generated the data. L.S. analyzed the data. L.S. and C.D.B. drafted the manuscript and interpreted the data. All the authors read and approved the manuscript.

---

**Supplementary Table S1:**
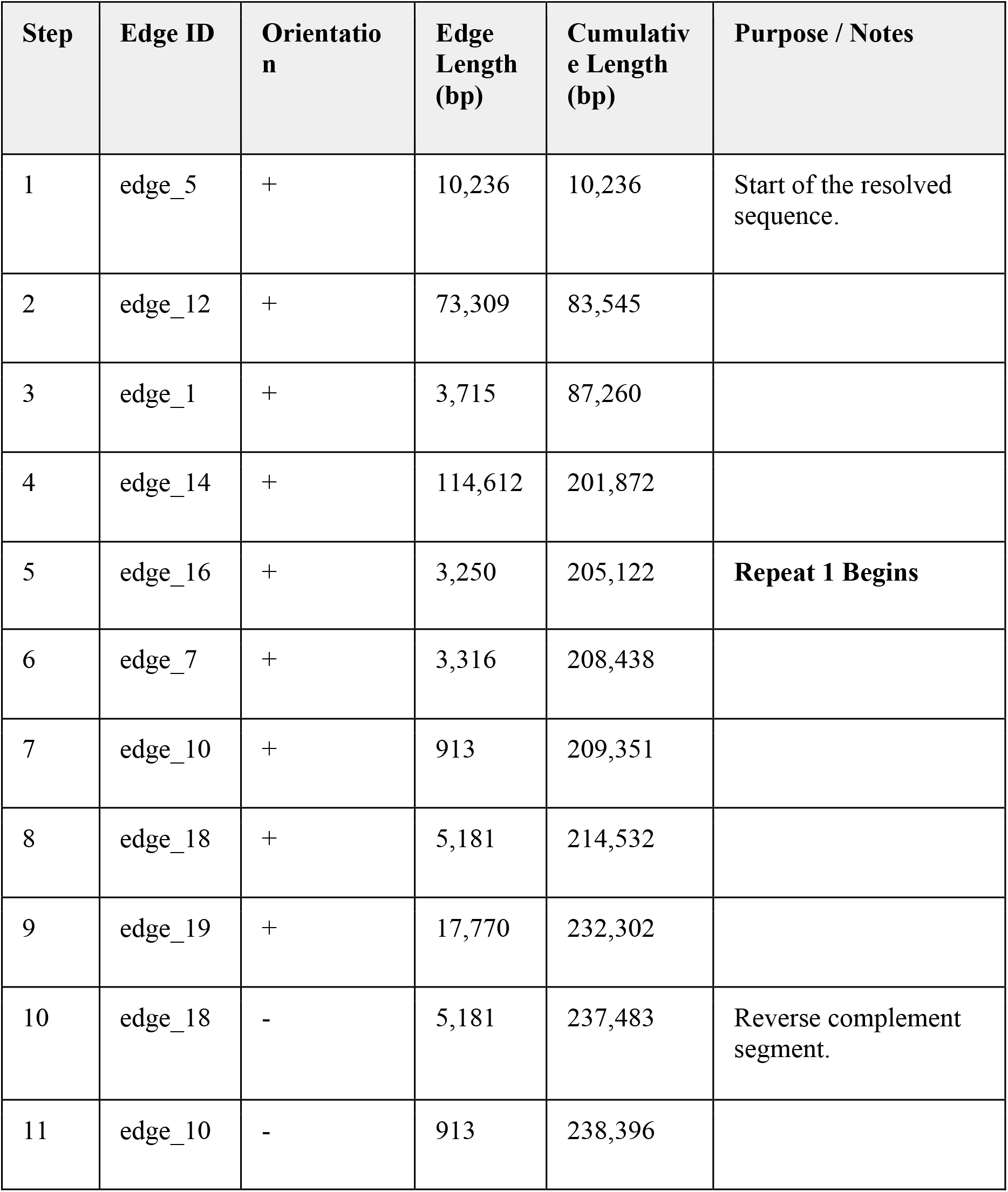
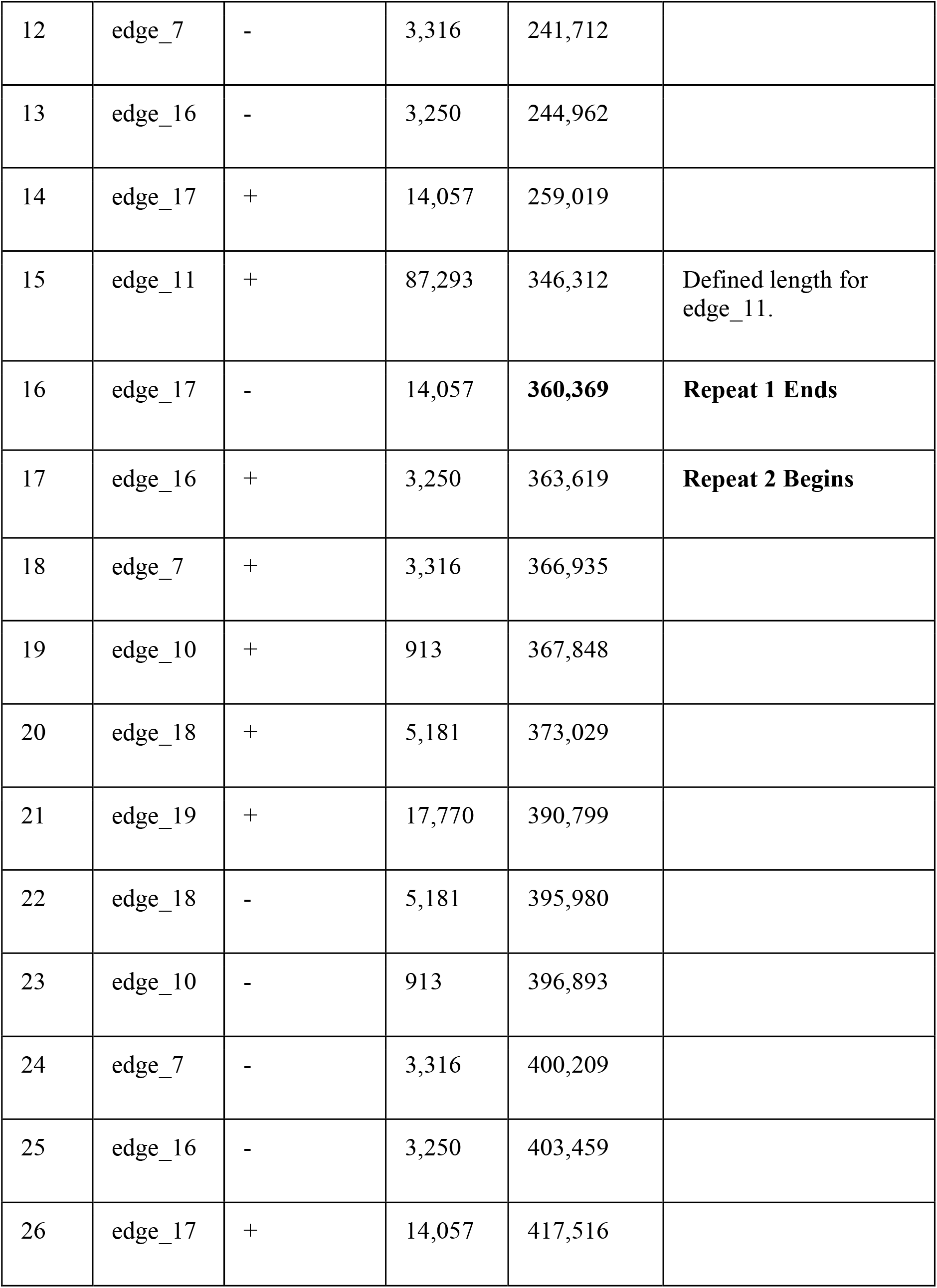
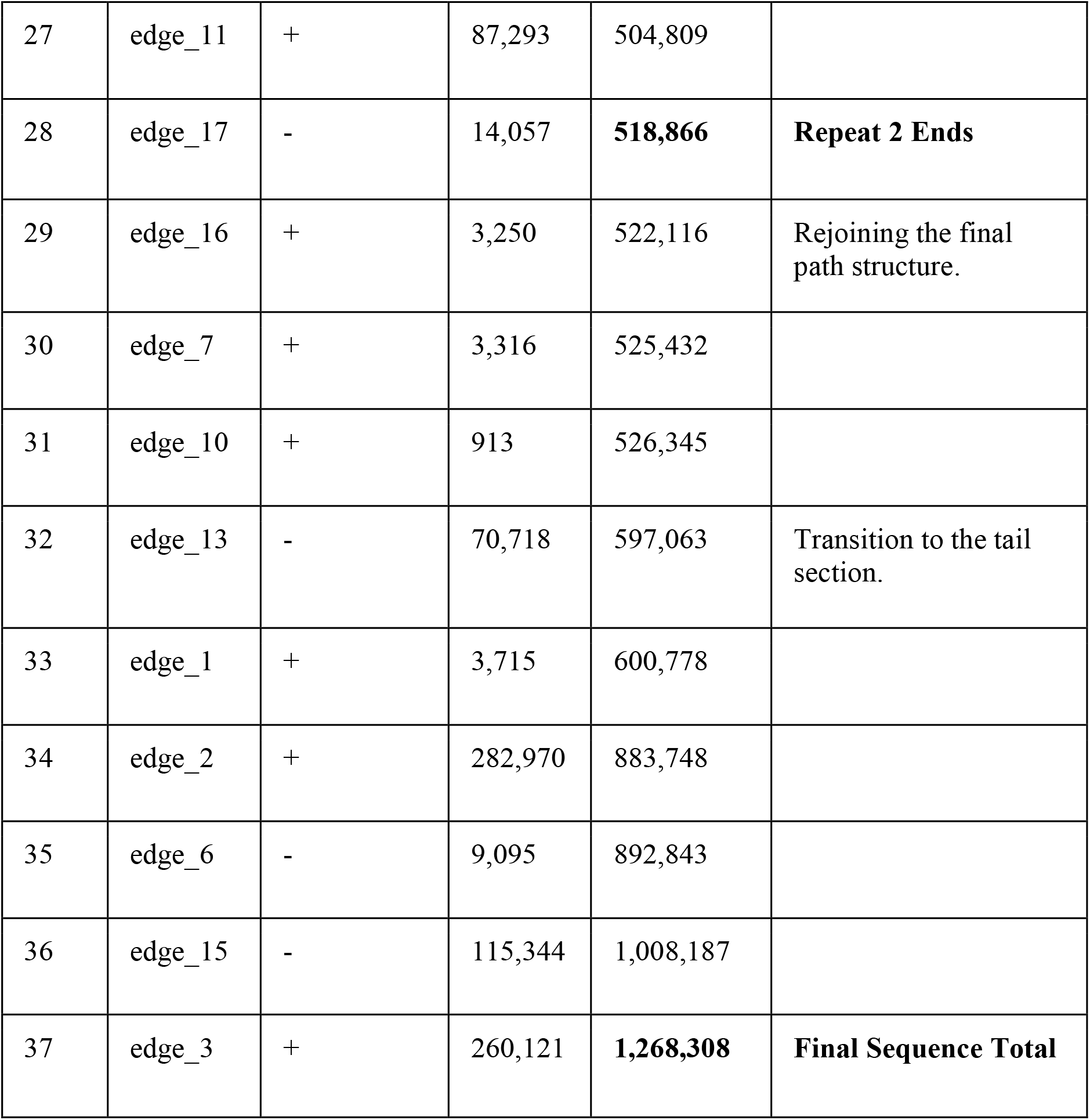
Manual resolution path of the *Euterpe precatoria* mitochondrial genome. Details the sequence of edges (contigs) and their respective orientations utilized for the manual resolution of the **Flye assembly graph** (derived from the *Cocos nucifera*-mapped HiFi reads). The path starts at the beginning of edge_5+ and concludes at the end of edge_3+, forming the **1**,**268**,**308 bp** linear master sequence. Edge_4, corresponding to a confirmed Nuclear Mitochondrial DNA segment (NUMT), was intentionally excluded from this path.

**Supplemental Figure 1.**
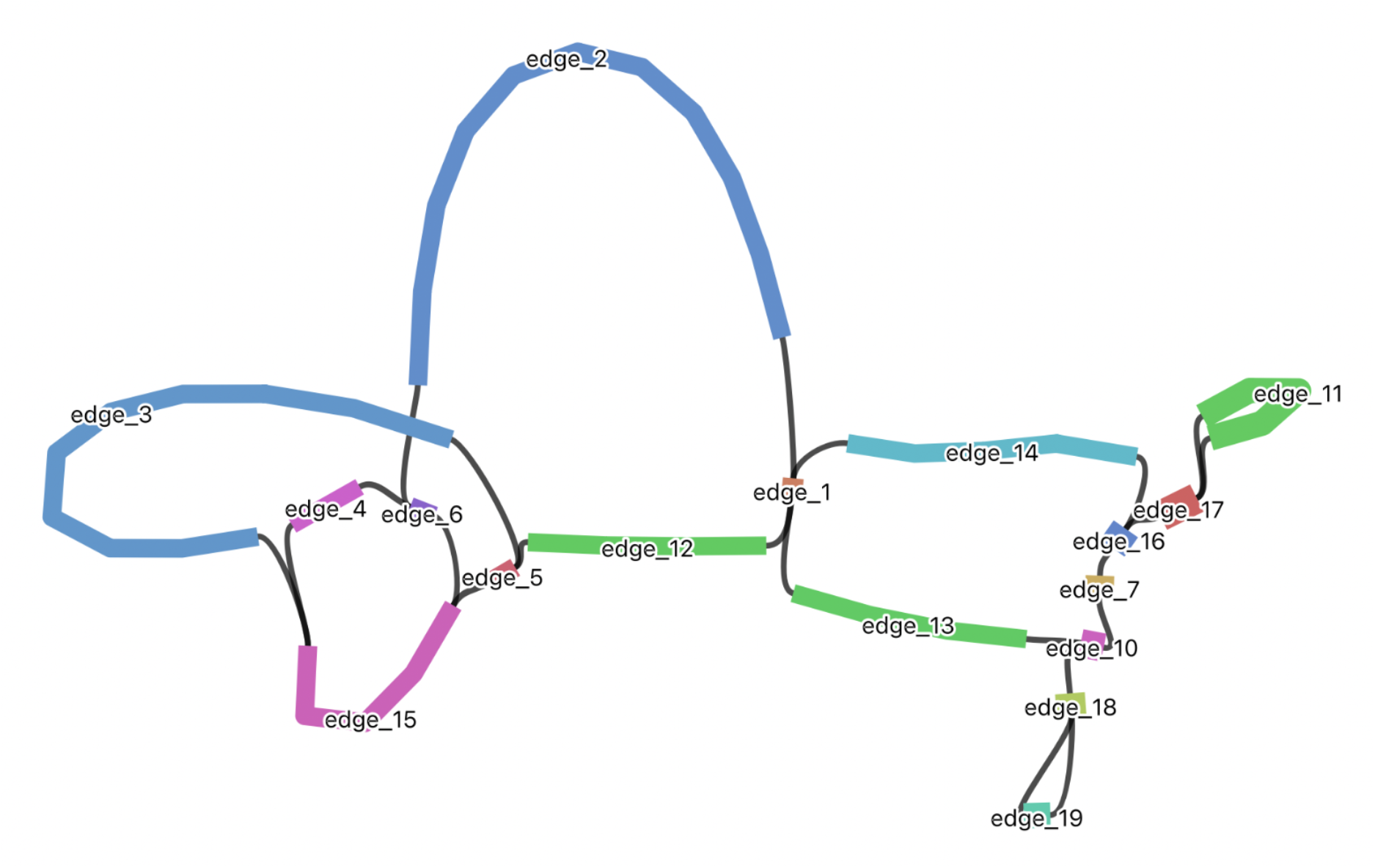
Visualization of the *Euterpe precatoria* mitochondrial genome assembly graph generated by Flye (version 2.9.1) and rendered in Bandage (version 0.8.1) with all edges marked with a numerical identifier.

