## Supplementary material for "A novel mitogenome for *Euterpe precatoria* (Arecaceae)": SI

Supplemental Figure 1. Visualization of the *Euterpe precatoria* mitochondrial genome assembly graph generated by Flye (version 2.9.1) and rendered in Bandage (version 0.8.1) with all edges marked with a numerical identifier.
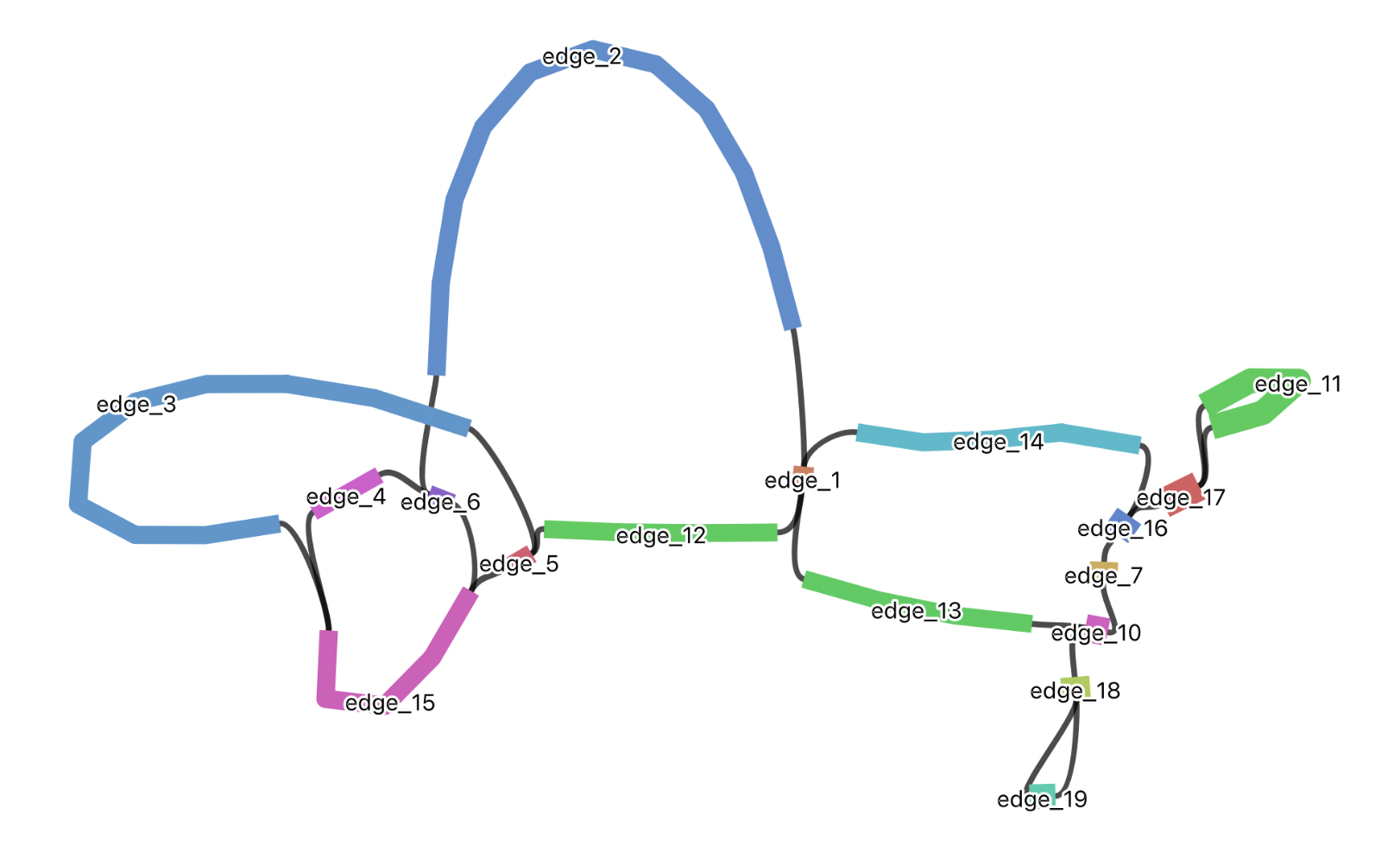

### **Supplementary Table S1: Manual resolution path of the *Euterpe precatoria* mitochondrial genome.** Details the sequence of edges (contigs) and their respective orientations utilized for the manual resolution of the **Flye assembly graph** (derived from the *Cocos nucifera*-mapped HiFi reads). The path starts at the beginning of edge_5+ and concludes at the end of edge_3+, forming the **1,268,308 bp** linear master sequence. Edge_4, corresponding to a confirmed Nuclear Mitochondrial DNA segment (NUMT), was intentionally excluded from this path.

| **Step** | **Edge ID** | **Orientation** | **Edge Length (bp)** | **Cumulative Length (bp)** | **Purpose / Notes** |
| --- | --- | --- | --- | --- | --- |
| 1 | edge_5 | + | 10,236 | 10,236 | Start of the resolved sequence. |
| 2 | edge_12 | + | 73,309 | 83,545 |  |
| 3 | edge_1 | + | 3,715 | 87,260 |  |
| 4 | edge_14 | + | 114,612 | 201,872 |  |
| 5 | edge_16 | + | 3,250 | 205,122 | **Repeat 1 Begins** |
| 6 | edge_7 | + | 3,316 | 208,438 |  |
| 7 | edge_10 | + | 913 | 209,351 |  |
| 8 | edge_18 | + | 5,181 | 214,532 |  |
| 9 | edge_19 | + | 17,770 | 232,302 |  |
| 10 | edge_18 | - | 5,181 | 237,483 | Reverse complement segment. |
| 11 | edge_10 | - | 913 | 238,396 |  |
| 12 | edge_7 | - | 3,316 | 241,712 |  |
| 13 | edge_16 | - | 3,250 | 244,962 |  |
| 14 | edge_17 | + | 14,057 | 259,019 |  |
| 15 | edge_11 | + | 87,293 | 346,312 | Defined length for edge_11. |
| 16 | edge_17 | - | 14,057 | **360,369** | **Repeat 1 Ends** |
| 17 | edge_16 | + | 3,250 | 363,619 | **Repeat 2 Begins** |
| 18 | edge_7 | + | 3,316 | 366,935 |  |
| 19 | edge_10 | + | 913 | 367,848 |  |
| 20 | edge_18 | + | 5,181 | 373,029 |  |
| 21 | edge_19 | + | 17,770 | 390,799 |  |
| 22 | edge_18 | - | 5,181 | 395,980 |  |
| 23 | edge_10 | - | 913 | 396,893 |  |
| 24 | edge_7 | - | 3,316 | 400,209 |  |
| 25 | edge_16 | - | 3,250 | 403,459 |  |
| 26 | edge_17 | + | 14,057 | 417,516 |  |
| 27 | edge_11 | + | 87,293 | 504,809 |  |
| 28 | edge_17 | - | 14,057 | **518,866** | **Repeat 2 Ends** |
| 29 | edge_16 | + | 3,250 | 522,116 | Rejoining the final path structure. |
| 30 | edge_7 | + | 3,316 | 525,432 |  |
| 31 | edge_10 | + | 913 | 526,345 |  |
| 32 | edge_13 | - | 70,718 | 597,063 | Transition to the tail section. |
| 33 | edge_1 | + | 3,715 | 600,778 |  |
| 34 | edge_2 | + | 282,970 | 883,748 |  |
| 35 | edge_6 | - | 9,095 | 892,843 |  |
| 36 | edge_15 | - | 115,344 | 1,008,187 |  |
| 37 | edge_3 | + | 260,121 | **1,268,308** | **Final Sequence Total** |
